# GLaMST: Grow Lineages along Minimum Spanning Tree for B Cell Receptor Sequencing Data

**DOI:** 10.1101/465476

**Authors:** Xingyu Yang, Christopher M. Tipton, Matthew C. Woodruff, Enlu Zhou, F. Eun-Hyung Lee, Inãki Sanz, Peng Qiu

## Abstract

B cell affinity maturation enables B cells to generate high-affinity antibodies. This process involves somatic hypermutation of B cell immunoglobulin receptor (BCR) genes and selection by their ability to bind antigens. Lineage trees are used to describe this microevolution of B cell immunoglobulin genes. In a lineage tree, each node is one BCR sequence that mutated from the germinal center and each directed edge represents a single base mutation, insertion or deletion. In BCR sequencing data, the observed data only contains a subset of BCR sequences in this microevolution process. Therefore, reconstructing the lineage tree from experimental data requires algorithms to build the tree based on partially observed tree nodes. We developed a new algorithm named Grow Lineages along Minimum Spanning Tree (GLaMST), which efficiently reconstruct the lineage tree given observed BCR sequences that correspond to a subset of the tree nodes. GLaMST constructs the minimum-spanning-tree (MST) to approximate the landscape of how observed BCR sequences are related, uses the MST to guide the interpolation of the closest unobserved sequence, updates the MST for the interpolation of additional unobserved sequences, and iterates until a full lineage tree is completed, where all observed sequences are connected by interpolated unobserved sequences and single base operations of mutations, insertions and deletions. Through comparison using simulated and real data, GLaMST outperforms existing algorithms in simulations with high rates of mutation, insertion and deletion, and generates lineage trees with smaller size and closer to ground truth according to tree features that highly correlated with selection pressure.

## 1 Introduction

To specifically recognize and respond to different pathogens, adaptive immune system relies on a diverse repertoire of B cell immunoglobulin receptors (BCR). Such a diverse repertoire comes from recombination, somatic hypermutation of immunoglobulin (Ig) gene segments, and selection by their ability to bind pathogens. This process will generate numerous different BCRs. To explore the dynamic process of BCR affinity maturation, researchers have applied high throughput sequencing [1, 2, 3] to examine BCR repertoires and to construct lineage trees of BCR sequences [4, 5].

In a BCR lineage tree, each tree node corresponds to one unique sequence, and each directed edge indicates the relationship between one sequence and its immediate ancestor, which are separated by one-base mutation, insertion or deletion. Given high throughput BCR sequencing data of a repertoire, the observed sequences correspond to some of the internal nodes and the leaf nodes of the BCR lineage tree, while many intermediate nodes are not observed due to the diversification and selection process the repertoire went through, as well as subsampling inherent to the assay. With these observed sequences, we can easily identify the root sequence of this tree by sequence alignment against known germline BCR segments in the genome [6]. To reconstruct the full lineage tree, we need to fill in the unobserved internal nodes and connect them to the observed nodes by direct edges. This process is similar to building the phylogenetic tree among species, except that only leaf nodes are observed in phylogenetic problem, whereas some internal nodes and the root nodes are also observed in this BCR lineage tree reconstruction problem. Popular methods in phylogenetic analysis includes maximum parsimony, maximum likelihood, and Bayesian methods. The maximum likelihood and Bayesian methods are usually computational demanding and require a decent amount of prior knowledge [7, 8], for example, the replacement rates and preference of mutation target under relatively selection pressure [9]. Such prior knowledge is relative limited in BCR lineage trees. Therefore, we decided to pursue the maximum parsimony idea to reconstruct the BCR lineage tree, which intends to reconstruct a tree as small as possible, which connects all the observed sequences with minimum number of mutation, insertion and deletion events.

Reconstruction of a phylogenetic tree using maximum parsimony method is known to be a NP-Complete problem [10, 11], which means we cannot guarantee a best solution in polynomial time. In terms of computational complexity, reconstruction of BCR lineage tree is very similar to phylogenetic tree. Although the root sequence and some of the internal nodes are known, it is still a NP-Complete problem [5]. To reconstruct BCR lineage trees from high throughput sequencing data, an algorithm named IgTree was previously developed, which is a heuristic procedure consisting of multiple components [5]. It first constructs a preliminary tree that only contains observed sequences based on multiple sequence alignment [12], then uses a complex scoring metric to gradually add internal node to complete the full tree, and finally scans the resulting tree to identify subtrees that can be reduced by reversion events. IgTree enabled efficient analysis of large BCR sequence datasets, and brought insights into various area of somatic-hypermutation-related biological processes including neutralization of HIV antibodies [13] and progression of follicular lymphoma [14]. Another algorithm for reconstructing BCR lineage trees is included in the TIgGER software package [15], which is a classical maximum parsimony method called “dnapars” in the PHYLIP library [16, 17]. dnapars almost always achieves smaller lineage trees compared to IgTree, but is considerably slower when analyzing large number of observed sequences.

Here, we present a novel algorithm, GLaMST, to reconstruct BCR lineage trees using the maximum parsimony criterion. GLaMST uses simple heuristics to Grow the Lineage trees along the Minimum Spanning Tree. In terms of the maximum parsimony criterion, GLaMST generates lineage trees with small size similar to dnapars, and outperforms dnapars for simulated datasets with insertions and deletions. In terms of the computational efficiency, GLaMST runs faster than IgTree. In this paper, we present the GLaMST algorithm, and evaluate its performance on both simulated and real data.

## 2 Methods

A formal description of the BCR lineage tree reconstruction is as follows. Given a set of observed BCR sequences and a root sequence, the maximum parsimony criterion would like to identify the minimum-sized directed tree structure with necessary intermediate sequences, where each directed edge in the tree represents a one-base operation (mutation, insertion and deletion), and all observed sequences are reachable from the root.

GLaMST reconstructs BCR lineage trees based on the minimum-spanning-tree (MST) [18, 19]. Figure 1 shows an outline of this algorithm. We first compute the pairwise edit-distances [20] of the observed sequences including the root node. These pairwise distances reflect the landscape of the observed sequences, in terms of their relative distances and directions with respect to the root node. The algorithm is initialized by considering the root sequence as the root node of the tree, the observed sequences as observed nodes, and no edges. We then grow the tree from the root node by adding directed edges and necessary intermediate nodes toward directions that are more populated by the observed sequences, and iteratively grow the tree until all the observed nodes are reachable from the root node. Observed nodes can either be internal nodes or leaf nodes of the tree, depending on whether they have descendants that are also observed nodes.

**Figure 1:**
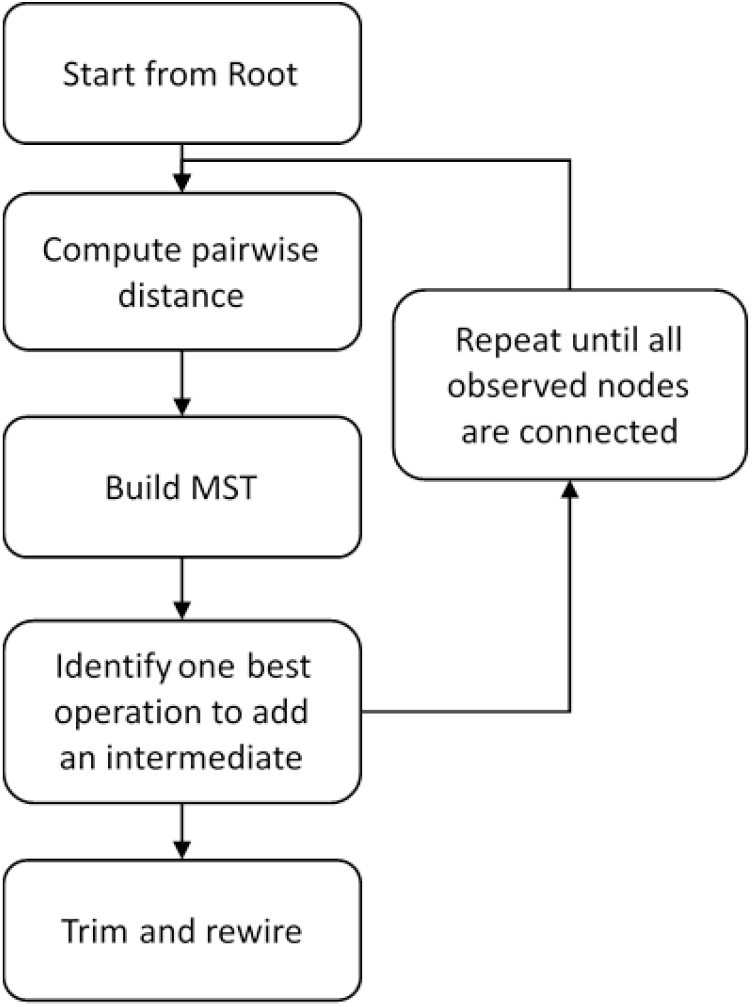
Overview of GLaMST.

### 2.1 Compute edit-distance

The edit-distance from one sequence to another sequence is the size of the minimal set of operations that can convert the first sequence into the second one [20]. In the context of DNA or RNA sequence alignment, one operation is mutation, insertion, or deletion of one base position. This distance metric is symmetric. We compute the pairwise edit-distances using the Wagner-Fischer algorithm, which is a dynamic programming algorithm with time complexity of *o*(*mn*), where *m* and *n* are the lengths of the two query sequences [21].

When computing the edit-distance from one sequence to another, we record the one-base operations in the minimal set. If there are multiple minimal sets, we record all operations in those sets, counting each unique operation only once. One example is shown in Figure 2. From sequence “ATCCCC” to “GCCCC”, the edit-distance is 2, because at least 2 one-base operations are needed to convert the first sequence to the second. As shown in Figure 2, there exist four paths of length 2 between the two sequences, and therefore, four possible sets of operations corresponding to the edit-distance. Out of the eight operations, four are unique (delete the 1st position, delete the 2nd position, mutate the 1st position to G, mutate the 2nd position to G). These four unique operations are recorded. The recorded operations reflect the “direction” from one sequence to the other, showing what operations can take the first sequence one step toward the second one. This direction information is useful in the next step to choose edges (operations) and intermediate nodes to grow the tree.

**Figure 2:**
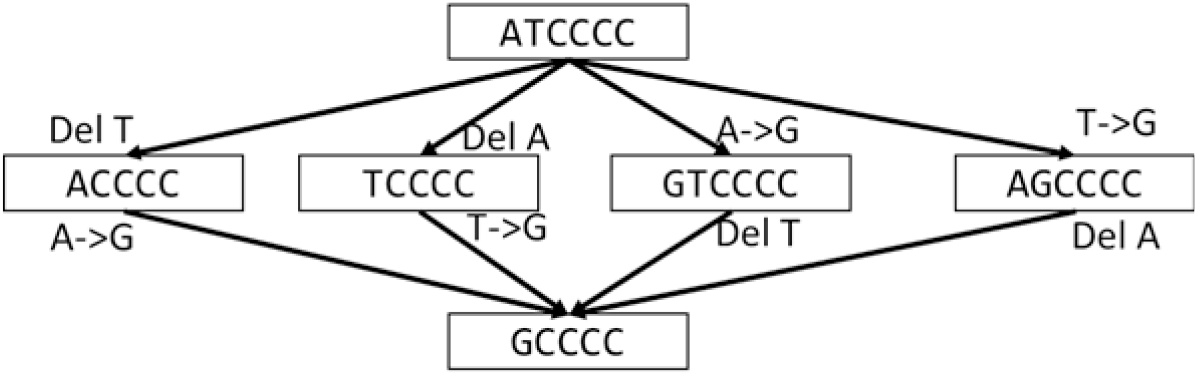
Example of edit-distance. The edit-distance between these two sequences is 2. There are four sets of operations corresponding to the edit-distance.

### 2.2 Initialize the lineage tree

We initialize GLaMST by treating the root sequence as the root node of the tree, and the observed sequence as other tree nodes. This initial structure does not contain any edges. The root node is considered as the reconstructed part of the lineage tree, whereas all other nodes are standing by, waiting to be brought into the reconstructed part of the lineage tree. Figure 3a shows an illustrative example of this initial structure and the distinction between the reconstructed part and the standby nodes.

**Figure 3:**
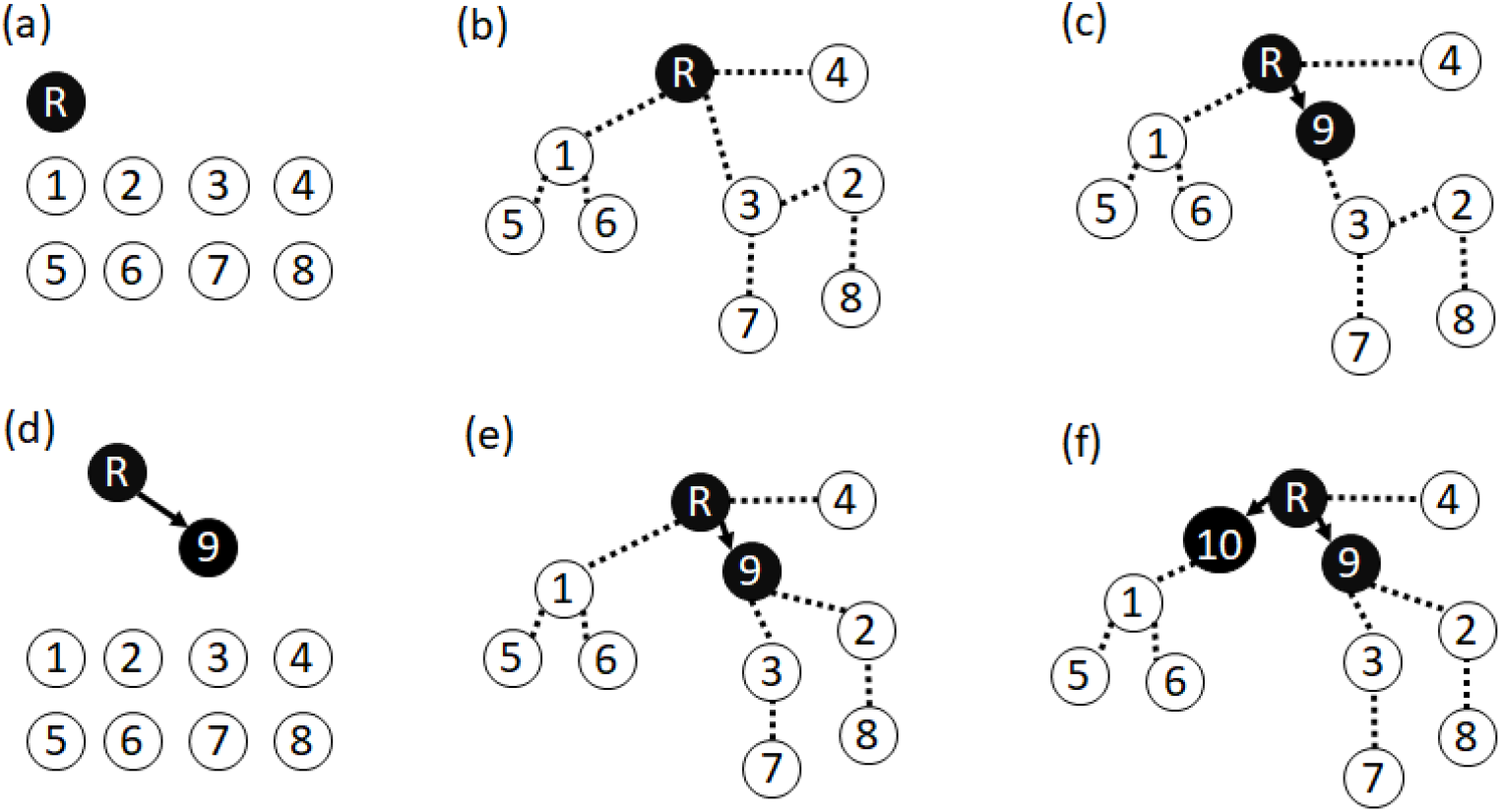
GLaMST tree construction process. This example shows the first two iterations of an illustrative example. In each visualization, black nodes and solid arrows represent the reconstructed part of the lineage tree. White nodes represent observed sequences that are standing by and waiting to be incorporated into the reconstructed part of the lineage tree. The dotted lines represent the MST, which guides the algorithm in growing the reconstructed part of the lineage tree. (a) GLaMST is initialized by treating the root node as the reconstructed part and all other observed nodes as standing by. (b) An MST is constructed based on the pairwise edit-distance. (c) GLaMST selects the most frequent origination-operation pair to create and insert an intermediate node and grow the reconstructed part. (d-f) The second iteration of the process to insert another node to the reconstructed part.

### 2.3 Iteratively grow the lineage tree

In the first iteration, GLaMST starts by building an MST using the pairwise edit-distances between all nodes. The MST is an undirected tree that connects all nodes with minimum total distances along its edges [19]. An illustrative example is shown in Figure 3b. Since the MST is undirected and the edit-distance associated to each edge is often larger than 1, edges of the MST are different from edges of the lineage tree we want to reconstruct. Figure 3b uses the dotted undirected lines to represent the MST.

The MST approximates the landscape of the observed sequences with respect to the reconstructed part of the lineage tree, grouping the standby nodes into clusters. For example, in Figure 3b, the observed sequences (standby nodes) are divided into three clusters, which locate in three “directions” from the root node. One cluster consists of nodes {2, 3, 7, 8} in the same direction from the root node, because they are relatively close to each other, and away from the root node and other nodes. There may exist one or several one-base operations that can take the root sequence one step closer to all those four nodes. Such operations are likely to generate intermediate nodes in the maximal parsimony lineage tree.

We then examine clusters of standby nodes originating from the same node in the reconstructed part of the lineage tree, which are all three clusters in Figure 3b attached to the root node. We take the sets of operations recorded when computing the edit-distances from the root node to the members in these clusters, and count the number of times each operation appears. If one operation appears four times, applying this operation to the root sequence will generate an intermediate sequence that is one step closer to four standby sequences. We choose the operation that appears the most number of times, and apply it to the root sequence to generate the first intermediate node to be added to the reconstructed part of the lineage tree. As shown in the illustrative example in Figure 3c, the reconstructed part now contains two nodes and one directed edge. The added node is typically along one branch of the MST, representing a common ancestor of the observed nodes in one cluster, but it is also possible that the added node is a common ancestor of multiple clusters.

The second iteration starts with the reconstructed part of the lineage tree and the standby nodes, as shown in Figure 3d. The previous MST is discarded, because the newly added node may cause the structure of the MST to change. In this iteration, we rebuild a new MST to connect the standby nodes to the reconstructed part of the lineage tree, as shown in Figure 3e. The new MST divides the standby nodes into four clusters. Two clusters originate from the root node, and two clusters originate from node 9. We examine the one-base operations for clusters attached to the same node in the reconstructed part of the tree (operations that take root node R to nodes {4} and {1, 5, 6}, and operations that take node 9 to nodes {2, 8} and {3, 7}), choose the origination-operation pair that appears the most number of times, and apply the operation to the origination node to generate an intermediate node to be added to the reconstructed part of the lineage tree. In the illustrative example in Figure 3f, the chosen origination-operation pair generated an intermediate node from the root node, which is one step closer toward the cluster {1, 5, 6}.

The subsequent iterations operate exactly the same as the second iteration. When selecting the most frequently appearing origination-operation pair, if there is a tie, we randomly choose one. When the new node suggested by the chosen origination-operation pair is identical to an observed node, the observed node is recruited into the reconstructed part of the lineage tree using a directed edge from the origination node to the observed node, and no intermediate node is added. The iteration continues until all observed nodes are recruited into the reconstructed part of the lineage tree.

### 2.4 Trim and rewire the lineage tree

The lineage tree reconstructed by this heuristic iterative procedure can be reduced by trimming off unnecessary branches. The iterative procedure may occasionally produce branches whose leaf nodes are not observed nodes, because the process of growing the tree is guided by the MST which only approximates the structure of the underlying lineage tree. Branches with unobserved nodes as leaves are unnecessary to explain the mutation process that gives rise to the observed nodes. To trim the lineage tree, we remove all unobserved intermediate and leaf nodes that do not have any observed nodes as descendants.

Another possible improvement is rewiring. In the reconstructed lineage tree, we can detach a subtree by removing the edge pointing to the root of the subtree, reattached it to some other node, and then trim the resulting tree. The trimming operation removes intermediate unobserved nodes right upstream of the removed edge. The reattaching operation introduces additional intermediate nodes if the edit-distance is larger than one between the subtree root and the node it reattaches to. It is possible that such a rewiring operation can reduce the size of the lineage tree. We consider all the observed nodes and branching nodes (i.e. out-degree larger than one) for rewiring. For each node under rewiring consideration, we try to rewire it to all possible nodes in the lineage tree, and examine whether we can reduce the tree size. If yes, we accept the rewiring operation, and examine the resulting lineage tree again for possible rewiring operations that may further reduce the tree size. We repeat this process until no rewiring operation can reduce the tree size.

## 3 Results

### 3.1 Reconstructed lineage trees using simulated data

To compare IgTree, dnapars and GLaMST, we generated simulated datasets using nine different simulation settings, varying the root sequence length and the relative probabilities of mutation, insertion, and deletion as shown in first column of Table 1. Following parameters in a previous simulation study [5], we generated 500 random lineage trees in each simulation setting, and subsampled the tree nodes to obtain the observed sequences. The overall tree size ranged from 20 to 80, and the number of observed nodes ranged from 2 to 24. Therefore, each simulated lineage tree contained 20~80 BCR sequences randomly generated by mutations, insertions or deletions of the root sequence, and the number of observed sequences ranged from 2 to 24. The simulated lineage trees served as the ground truth that we would like to recover using the algorithms. Table 1 shows the comparison of size of lineage trees reconstructed by three different algorithms in all nine simulated datasets.

**Table 1:** Comparison of tree size based on simulated data.

| Simulation | Seq Length | Mutation Distribution<br>(mutation, insertion, deletion) | IgTree |  |  |  | dnapars |  |  |  | GLaMST |  |  |  |
| --- | --- | --- | --- | --- | --- | --- | --- | --- | --- | --- | --- | --- | --- | --- |
|  |  |  | Larger | Same | Smaller | Success | Larger | Same | Smaller | Success | Larger | Same | Smaller | Success |
| 1 | 300 | (1.00, 0.00, 0.00) | 10 | 469 | 21 | 98% | 0 | 480 | 20 | 100% | 6 | 473 | 21 | 99% |
| 2 | 300 | (0.98, 0.01, 0.01) | 12 | 468 | 20 | 98% | 3 | 477 | 20 | 99% | 7 | 472 | 21 | 99% |
| 3 | 300 | (0.90, 0.05, 0.05) | 54 | 424 | 22 | 89% | 34 | 444 | 22 | 93% | 11 | 464 | 25 | 98% |
| 4 | 80 | (1.00, 0.00, 0.00) | 39 | 390 | 71 | 92% | 1 | 420 | 79 | 100% | 38 | 391 | 71 | 92% |
| 5 | 80 | (0.98, 0.01, 0.01) | 71 | 373 | 56 | 86% | 20 | 416 | 64 | 96% | 28 | 414 | 58 | 94% |
| 6 | 80 | (0.90, 0.05, 0.05) | 130 | 325 | 45 | 74% | 103 | 345 | 52 | 79% | 26 | 412 | 62 | 95% |
| 7 | 20 | (1.00, 0.00, 0.00) | 77 | 234 | 189 | 85% | 8 | 241 | 251 | 98% | 46 | 251 | 203 | 91% |
| 8 | 20 | (0.98, 0.01, 0.01) | 116 | 201 | 183 | 77% | 54 | 221 | 225 | 89% | 35 | 236 | 229 | 93% |
| 9 | 20 | (0.90, 0.05, 0.05) | 278 | 129 | 93 | 44% | 234 | 153 | 113 | 53% | 47 | 255 | 198 | 91% |

As shown in Table 1, in the first simulation setting where the sequence length was 300 and probabilities of insertion and deletion were 0, all three methods performed well. We recorded how many times the size of reconstructed tree is smaller, equal, or larger than the simulated ground truth. A reconstructed tree larger than the simulated ground truth was undesirable, because the reconstruction failed to achieve the goal of maximum parsimony. A reconstructed tree could occasionally be smaller than the simulated ground truth. This was because the observed sequences represented only a subset of the simulated mutation process, and it was possible to explain such partial observations by trees smaller than the simulated mutation process. We considered it a success if the reconstructed tree was of smaller or equal size compared to the simulated ground truth. In the first simulation setting, the success rates of all three algorithms were >98%. This was mainly because the first simulation setting was relatively simple: given a sequence length of 300 and the underlying tree size of 20~80, the simulated mutations seldom occurred more than once at the same position.

Comparing the first three simulation settings with the same root sequence length but different proportions of mutations, insertions and deletions, we can see that the performance of all three algorithms decreased with increased insertions and deletions. The same trend was seen in simulation settings with shorter sequence length, showing that the presence of insertions and deletions made the maximum parsimony lineage tree reconstruction problem more challenging.

In the first set of three simulations, the mutations, insertions and deletions are uniformly distributed in simulated sequences of length 300. In reality, the frequency of mutations is not uniform across the BCR. For example, the V(D)J region of the BCR can be partitioned into framework regions (FWRs) which typically have lower observed mutation frequencies, and complementarity determining regions (CDRs) which have higher observed mutation frequencies [6]. To examine a simpler situation that has the flavor of variable mutation rate, we created two subsequent sets of simulations with short sequence lengths of 80 and 20, respectively. These simulations were equivalent to simulating the length 300 sequences while constraining the mutations to only a sub-region of length either 80 or 20. These simulation settings with shorter sequence length were progressively more difficult, because the simulated mutations, insertions and deletions in shorter sequences were more likely to occur multiple times at the same position, which represented higher mutation rates. Table 1 shows that the reconstruction performance of all three algorithms consistently decreased with higher mutation rates.

Overall, in terms of tree size shown in Table 1, GLaMST and dnapars consistently outperformed IgTree in all nine simulation settings. In simulation settings without insertions and deletions, dnapars outperformed GLaMST by a relatively small margin. However, in simulation settings with insertions and deletions, GLaMST achieved the significantly better performance than dnapars, especially in the last and most challenging simulation setting.

In addition to tree size, we also compared the three algorithms based on tree features defined in the MTree program for lineage tree measurement [22, 23], especially features that are highly correlated with selection pressures [23]. We considered 12 features listed in Table 2. For each simulated lineage tree, we computed the difference between the tree features of the simulation ground truth and the tree features of the lineage trees reconstructed by all three algorithms, and then normalized the differences by dividing by the range of corresponding tree features in the simulation ground truth. The average differences for the simulation settings and algorithms are compared in Figure 4. For these 12 tree features, GLaMST achieved differences either smaller than or comparable to the other two algorithms, meaning that the lineage trees constructed by GLaMST were more similar to the simulation ground truth compared to the other two algorithms, especially in the most challenging simulation setting.

**Table 2:** Comparison of 12 tree features based on real data.

|  | Description | IgTree | dnapars | GLaMST |
| --- | --- | --- | --- | --- |
| Tree Size | Total number of tree nodes | 1396 | 1199 | 1190 |
| OD-Root | Out-degree of the root of the tree | 1 | 1 | 1 |
| OD-Avg | Average out-degree of all tree nodes | 0.999 | 0.999 | 0.999 |
| OD-Ratio | Ratio between OD-Root and OD-Avg | 1.001 | 1.001 | 1.001 |
| T | Maximum depth of the tree leaves | 62 | 147 | 130 |
| PL-Min | Minimum distance from root to any leaf | 12 | 38 | 36 |
| PL-Avg | Average distance from root to all nodes | 30.164 | 84.624 | 79.821 |
| DRSN-Min | Minimum distance from root to any branching node | 2 | 36 | 34 |
| DLFSN-Min | Minimum distance from first branching node to leaves | 10 | 2 | 2 |
| DLFSN-Avg | Average distance from first branching node to leaves | 28.164 | 48.624 | 45.821 |
| DASN-Min | Minimum distance between branching nodes | 1 | 1 | 1 |
| DASN-Avg | Average distance between branching nodes | 2.903 | 2.590 | 2.216 |

**Figure 4:**
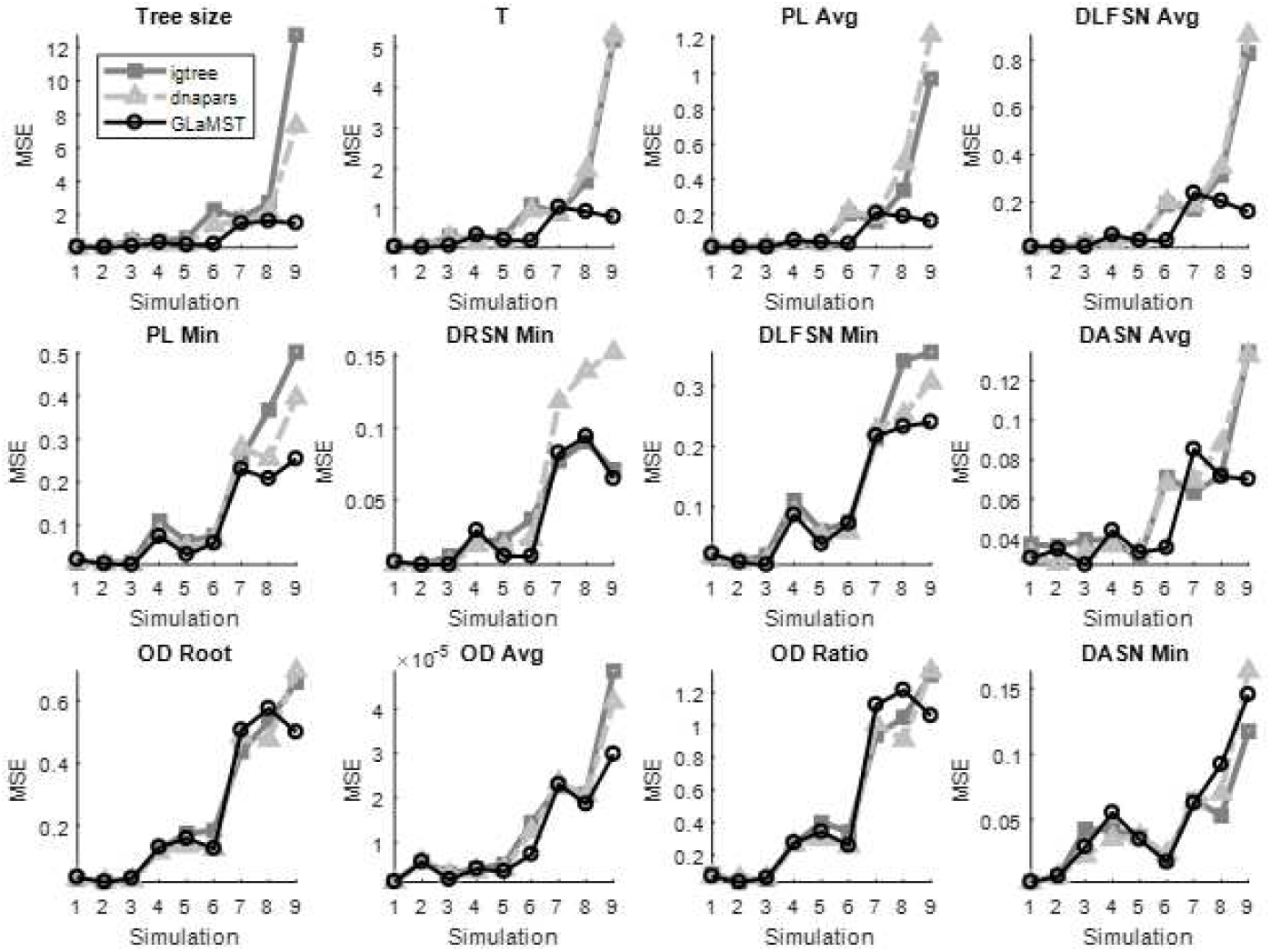
Comparison based on tree features in simulated data. These figures compare GLaMST, IgTree and dnapars using 12 tree features. Descriptions of these features are listed in Table 2. Each panel corresponds to one tree feature, x-axis represents the nine simulation settings and y-axis is the mean square error between reconstructed trees and simulated ground truth. Tree features in the first row suggest that GLaMST is significantly better than the other two methods. The second row shows tree features where GLaMST is moderately outperforms the other methods. The third row shows tree features where the three algorithms show similar performance.

### 3.2 Reconstructed lineage trees using BCR sequence data

In order to produce a biological dataset for testing, peripheral blood mononuclear cells (PBMC) were collected from an individual afflicted with Pemphigus Vulgaris. These cells were sorted via fluorescence-activated cell sorting (FACS) into 4 major B cell populations: Naive (CD19+IgD+CD27-), Switched memory (CD19+IgD-CD27+), Double negative (CD19+IgD-CD27-), and Plasmablasts (CD19+/-CD27++CD38++). RNA was extracted and immunoglobulin heavy chain transcripts were amplified using RT-PCR with primers located in the framework region 1 for VH families 1-7 and constant regions for IgM, IgG, and IgA. The amplicons were subsequently sequenced on an Illumina MiSeq using 300bp x2 paired end reads. Reads were processed using our IgSeq pipeline as described in [24], where sequences from all populations were quality filtered, joined, and divided into clones based on same V gene usage, same J gene usage, identical CDR3 length, and 85% CDR3 similarity. Therefore, the data for each individual clone consisted of sequences sharing the same V gene, the same J gene, and the same CDR3 length with 85% sequence similarity in the CDR3 region. Data for individual clones were then used as separate datasets for further testing of GLaMST, with the largest clone being illustrated here.

In the dataset corresponding to the largest clone, the lengths of the observed sequences were around 320, and the total number of observed sequences was 684. The root node sequence was derived from the most likely germline IgHV sequence as identified using IMGT/HighV-QUEST (http://www.IMGT.org), and trimmed to match the forward primer location and beginning of the CDR3 [25]. The root sequence had a 48-base segment at the five-prime end that did not exist in many of the observed sequences, as shown in Figure 5a, which visualized a few selected observed sequences using multiple alignment [12]. Therefore, a fixed size gap existed between root and many of the observed sequences. This gap was because there were two forward primer locations used in the experimental setup that generated the data. This gap can be removed by further trimming the root sequence and some of the observed sequences according to the location of the primer that was relatively downstream. However, even without such further preprocessing, successful lineage tree reconstruction algorithms should be able to recognize this gap as a common segment of deletion between the observed sequences and the root. Therefore, we decided to keep the root sequence as is and evaluate whether tree reconstruction algorithms could identify the common deletion shared by the observed sequences.

**Figure 5:**
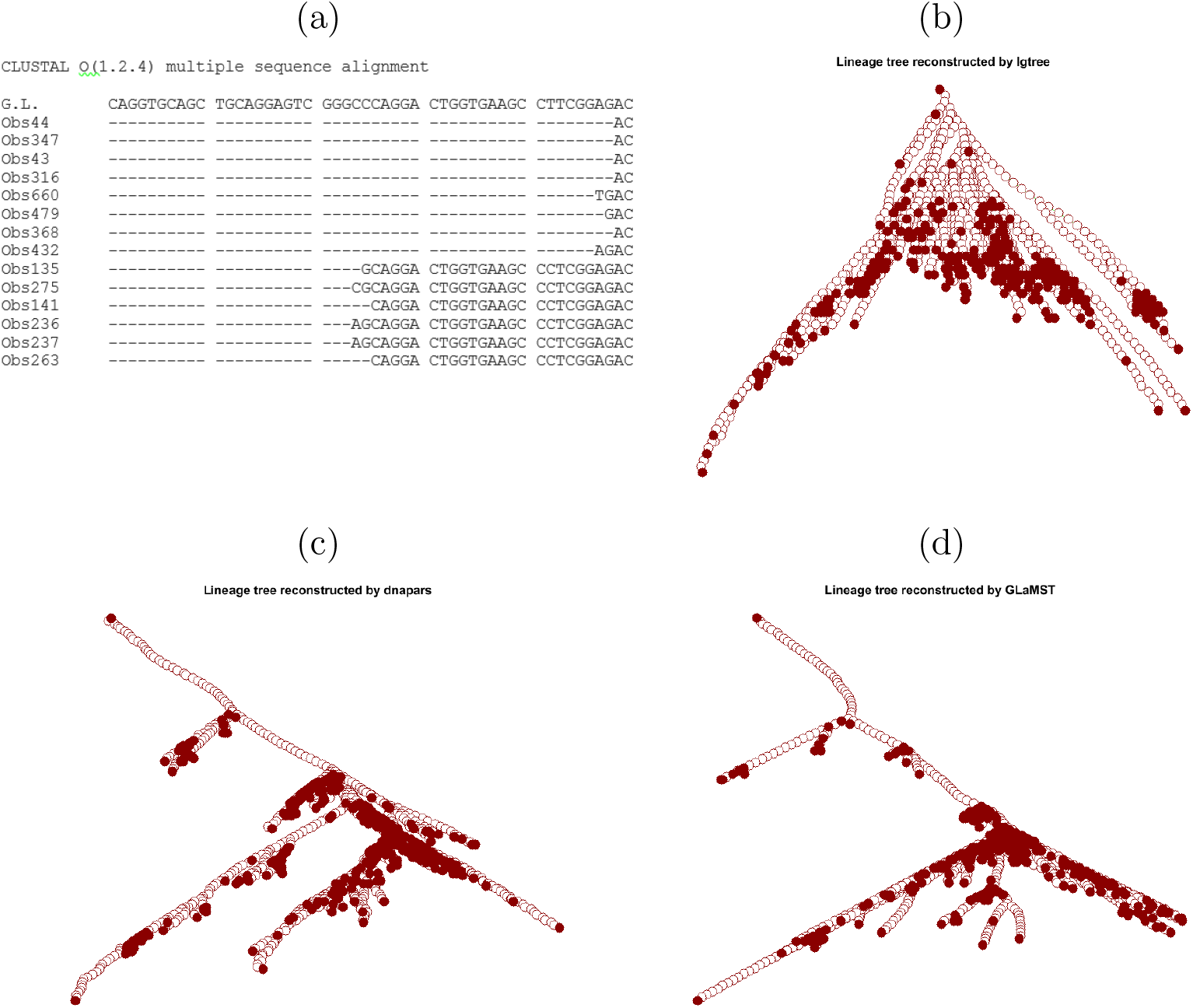
Comparison based on real data. (a) Multiple sequence alignment of the root node and 14 selected observed nodes. This sequence alignment shows a fixed gap between root node and observed nodes. (b) Lineage tree constructed by Igtree. Square represents the root node. The solid nodes are the observed sequences, while the white/empty nodes are intermediate nodes. (c) Lineage tree constructed by dnapars. (d) Lineage tree constructed by GLaMST.

The lineage tree reconstructed by IgTree was of size 1396 (Figure 5b). dnapars generated a lineage tree which has 1199 nodes (Figure 5c). The lineage tree reconstructed by GLaMST contained 1190 nodes (Figure 5d). Therefore, GLaMST and dnapars significantly outperformed IgTree in achieving the maximum parsimony criterion. From Figures 5c and 5d, we can see that the trees reconstructed by GLaMST and dnapars shared similar topology, with a long chain of intermediate nodes off of the root before the first branching point. This chain corresponded to the 48-base segment in the root sequence which did not exist in the observed sequences. In contrast, Figure 5b shows that IgTree failed to recognize this common difference between the observed sequences and the root, generating a tree wider and shallower tree with much larger size compared to GLaMST and dnapars. We also computed the 12 tree features in Table 2, which confirmed that the topology of the GlaMST and dnapars results were similar to each other, but very different from that of IgTree.

This dataset contained 684 observed sequences, which was much larger than the simulated data, and therefore, enabled us to compare the running time of the three algorithms. These algorithms were compared on a desktop computer with Intel i7-3770 processor at 3.40GHz and 32GM memory. dnapars spent roughly 5 days to reconstruct a lineage tree for this dataset. IgTree took 20 minutes, while GLaMST took only 16 minutes. Although dnapars and GLaMST reconstructed similar tree structures, GLaMST is significant more efficient computationally.

## 4 Conclusion

High throughput sequencing of BCR repertoire analysis motivated the computational question of reconstructing lineage trees based on partially observed tree nodes. We developed the GLaMST algorithm to reconstruct lineage trees with maximum parsimony. As suggested by its name, GLaMST grows lineage trees along the minimum spanning tree, which is a simple and efficient heuristic algorithm. Using both simulated and real data, we demonstrated that GLaMST is more effective in achieving maximum parsimony compared two existing algorithms. GLaMST is also computationally more efficient, enabling its application in analysis of large BCR sequencing datasets. Integrating GLaMST into existing BCR sequencing analysis frameworks can lead to significant improves in the lineage tree reconstruction aspect of the analysis.

## 5 Availability

Source code available at https://github.com/xysheep/GLaMST, and example demonstrations available at https://xysheep.github.io/GLaMST.

## Acknowledgement

This work was partially supported by funding from the National Science Foundation (CCF1552784). P.Q. is an ISAC Marylou Ingram Scholar.

## References

[1] J. J. Calis and B. R. Rosenberg. Characterizing immune repertoires by high throughput sequencing: strategies and applications. Trends Immunol., 35(12):581–590, Oct 2014.

[2] L. He, D. Sok, P. Azadnia, J. Hsueh, E. Landais, M. Simek, W. C. Koff, P. Poignard, D. R. Burton, and J. Zhu. Toward a more accurate view of human B-cell repertoire by next-generation sequencing, unbiased repertoire capture and single-molecule barcoding. Sci Rep, 4:6778, Oct 2014.

[3] G. Georgiou, G. C. Ippolito, J. Beausang, C. E. Busse, H. Wardemann, and S. R. Quake. The promise and challenge of high-throughput sequencing of the antibody repertoire. Nat. Biotechnol., 32(2):158–168, Feb 2014.

[4] D. K. Dunn-Walters, A. Belelovsky, H. Edelman, M. Banerjee, and R. Mehr. The dynamics of germinal centre selection as measured by graph-theoretical analysis of mutational lineage trees. Dev. Immunol., 9(4):233–243, Dec 2002.

[5] M. Barak, N. S. Zuckerman, H. Edelman, R. Unger, and R. Mehr. IgTree: creating Immunoglobulin variable region gene lineage trees. J. Immunol. Methods, 338(1-2):67–74, Sep 2008.

[6] G. Yaari and S. H. Kleinstein. Practical guidelines for B-cell receptor repertoire sequencing analysis. Genome Med, 7:121, Nov 2015.

[7] Z. Yang and B. Rannala. Molecular phylogenetics: principles and practice. Nat. Rev. Genet., 13(5):303–314, Mar 2012.

[8] A. V. Werhli and D. Husmeier. Gene regulatory network reconstruction by Bayesian integration of prior knowledge and/or different experimental conditions. J Bioinform Comput Biol, 6(3):543–572, Jun 2008.

[9] S. Whelan, P. Lio, and N. Goldman. Molecular phylogenetics: state-of-the-art methods for looking into the past. Trends Genet., 17(5):262–272, May 2001.

[10] W. H. Day, D. S. Johnson, and D. Sankoff. The computational complexity of inferring rooted phylogenies by parsimony. Mathematical Biosciences, 81(1):33–42, 1986.

[11] B. Chor and T. Tuller. Maximum Likelihood of Evolutionary Trees Is Hard, pages 296–310. Springer Berlin Heidelberg, 2005.

[12] J. D. Thompson, D. G. Higgins, and T. J. Gibson. CLUSTAL W: improving the sensitivity of progressive multiple sequence alignment through sequence weighting, position-specific gap penalties and weight matrix choice. Nucleic Acids Res., 22(22):4673–4680, Nov 1994.

[13] D. Sok, U. Laserson, J. Laserson, Y. Liu, F. Vigneault, J. P. Julien, B. Briney, A. Ramos, K. F. Saye, K. Le, A. Mahan, S. Wang, M. Kardar, G. Yaari, L. M. Walker, B. B. Simen, E. P. St John, P. Y. Chan-Hui, K. Swiderek, S. H. Kleinstein, S. H. Kleinstein, G. Alter, M. S. Seaman, A. K. Chakraborty, D. Koller, I. A. Wilson, G. M. Church, D. R. Burton, and P. Poignard. The effects of somatic hypermutation on neutralization and binding in the PGT121 family of broadly neutralizing HIV antibodies. PLoS Pathog., 9(11):e1003754, 2013.

[14] M. R. Green, A. J. Gentles, R. V. Nair, J. M. Irish, S. Kihira, C. L. Liu, I. Kela, E. S. Hopmans, J. H. Myklebust, H. Ji, S. K. Plevritis, R. Levy, and A. A. Alizadeh. Hierarchy in somatic mutations arising during genomic evolution and progression of follicular lymphoma. Blood, 121(9):1604–1611, Feb 2013.

[15] N. T. Gupta, J. A. Vander Heiden, M. Uduman, D. Gadala-Maria, G. Yaari, and S. H. Kleinstein. Change-O: a toolkit for analyzing large-scale B cell immunoglobulin repertoire sequencing data. Bioinformatics, 31(20):3356–3358, Oct 2015.

[16] J. Felsenstein. PHYLIP - phylogeny inference package (version 3.2). Cladistics, 5:164–166, 1989.

[17] J. N. Stern, G. Yaari, J. A. Vander Heiden, G. Church, W. F. Donahue, R. Q. Hintzen, A. J. Huttner, J. D. Laman, R. M. Nagra, A. Nylander, D. Pitt, S. Ramanan, B. A. Siddiqui, F. Vigneault, S. H. Kleinstein, D. A. Hafler, and K. C. O’Connor. B cells populating the multiple sclerosis brain mature in the draining cervical lymph nodes. Sci Transl Med, 6(248):248ra107, Aug 2014.

[18] J. B. Kruskal. On the shortest spanning subtree of a graph and the traveling salesman problem. Proceedings of the American Mathematical Society, 7(1):48–50, February 1956.

[19] P. Qiu, E. F. Simonds, S. C. Bendall, K. D. Gibbs, R. V. Bruggner, M. D. Linderman, K. Sachs, G. P. Nolan, and S. K. Plevritis. Extracting a cellular hierarchy from high-dimensional cytometry data with SPADE. Nat. Biotechnol., 29(10):886–891, Oct 2011.

[20] R. C. Prim. Shortest connection networks and some generalizations. Bell System Technology Journal, 36:1389–1401, 1957.

[21] R. A. Wagner and M. J. Fischer. The string-to-string correction problem. J. ACM, 21(1):168–173, January 1974.

[22] D. K. Dunn-Walters, H. Edelman, and R. Mehr. Immune system learning and memory quantified by graphical analysis of B-lymphocyte phylogenetic trees. BioSystems, 76(1-3):141–155, 2004.

[23] M. Uduman, M. J. Shlomchik, F. Vigneault, G. M. Church, and S. H. Kleinstein. Integrating B cell lineage information into statistical tests for detecting selection in Ig sequences. J. Immunol., 192(3):867–874, Feb 2014.

[24] C. M. Tipton, C. F. Fucile, J. Darce, A. Chida, T. Ichikawa, I. Gregoretti, S. Schieferl, J. Hom, S. Jenks, R. J. Feldman, R. Mehr, C. Wei, F. E. Lee, W. C. Cheung, A. F. Rosenberg, and I. Sanz. Diversity, cellular origin and autoreactivity of antibody-secreting cell population expansions in acute systemic lupus erythematosus. Nat. Immunol., 16(7):755–765, Jul 2015.

[25] Eltaf Alamyar, Vronique Giudicelli, Shuo Li, Patrice Duroux, and Marie-Paule Lefranc. Imgt/highv-quest: the imgt web portal for immunoglobulin (ig) or antibody and t cell receptor (tr) analysis from ngs high throughput and deep sequencing. 8:26, 01 2012.

